# Noise induced bimodality in genetic circuits with monostable positive feedback

**DOI:** 10.1101/464297

**Authors:** Pavol Bokes, Abhyudai Singh

## Abstract

The expression of individual genes can be maintained through positive feedback loop mechanisms. If genes are expressed in bursts, then feedback either affects the frequency with which bursts occur or their size. Here we use a tractable hybrid modelling framework to evaluate how noncooperative positive feedback in burst frequency or burst size impacts the protein-level distribution. We confirm the results of previous studies that noncooperative positive feedback in burst frequency can support bimodal distributions. Intriguingly, bimodal distributions are unavailable in the case of feedback in burst size in the hybrid framework. However, kinetic Monte Carlo simulations of a full discrete model show that bimodality can reappear due to low-copy number effects. The two types of feedbacks lead to dramatically different values of protein mean and noise. We show that small values of leakage imply a small protein mean for feedback in burst frequency but not necessarily for feedback in burst size. We also show that protein noise decreases in response to gene activation if feedback is in burst frequency but there is a transient noise amplification if feedback acts on burst size. Our results suggest that feedback in burst size and feedback in burst frequency may play fundamentally different roles in maintaining and controlling stochastic gene expression.

## I. Introduction

The ultimate product of gene expression, the protein, is often synthesised in bursts of many molecule copies [1], [2]. Burst-like synthesis of protein is thought to be a substantive contributor towards the total gene-expression noise [3], [4]. Noise in gene expression drives variability in isogenic cell populations as well as temporal fluctuations in the phenotypes of single cells [5], [6].

Positive feedback loop provides a mechanism by which a protein can maintain its gene expression [7]. Positive feedback can support two stable states of protein expression in the deterministic setting and bimodal protein distributions in the stochastic setting [8]. Deterministic analysis shows that bistable behaviour requires cooperativity in feedback [9]. Stochastic studies indicate that bimodal distributions are available without cooperativity [10]–[17]. Random fluctuations in protein level can lead to extinction in a stochastic setting [18]. An ultimate extinction can be avoided by considering the effects of leakage — a small rate of spontaneous gene expression which occurs independently of the feedback.

In this paper we study the control of burst frequency and burst size through a noncooperative positive feedback with leakage. We adapt a minimalistic modelling framework [19]–[21] in which protein decays deterministically but is produced in discontinuous bursts (Fig 1, left). The chosen framework belongs to a wider family of piecewise deterministic [22], [23] or hybrid models [24], [25]. Regulation of burst frequency (Fig 1, right top), and partly also burst size (Fig 1, right bottom), have previously been examined using the aforementioned modelling framework [26]–[29]. In the context of burst-size regulation, two alternative versions have been proposed depending on the inclusion or omission of a so-called infinitesimal delay [30], [31]. Here we shall focus specifically on the undelayed case in the sense of [30], [31].

**Fig. 1.**
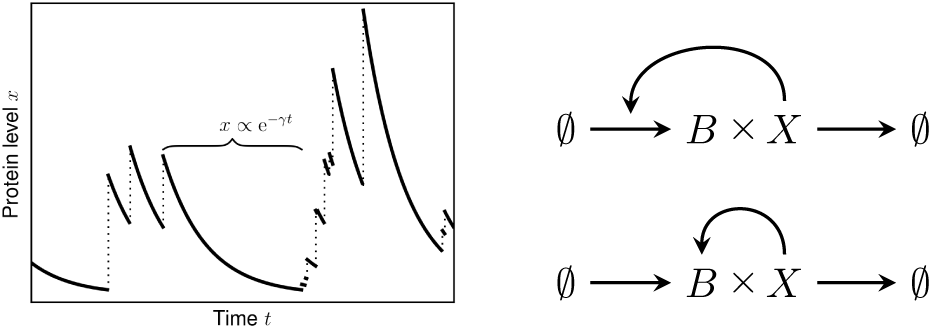
*Left:* Sample path of protein dynamics. In between bursts, protein level decays exponentially with rate constant *γ* (solid lines). Bursts occur randomly in time and lead to positive discontinuous jumps in protein level (dotted vertical lines). *Right top:* Feedback in burst frequency: protein X, which is produced in bursts of size B as well as being removed from the system, maintains the frequency of burst occurrence. *Right bottom:* Feedback in burst size: same as above, but here protein X maintains the size of its bursts.

Section II introduces the model and its master equation, explaining the differences between feedback in burst frequency and burst size. Section III recapitulates analytic results for feedback in burst frequency, some of which are known from previous studies. Section IV provides analogous analytic results for feedback in burst size. Section V presents discrete reaction systems that incorporate feedback in burst frequency or burst size. Section VI applies the theoretical analysis of the previous sections to gain a thorough understanding of the most distinctive properties of the feedback models. Section VII concludes the paper.

## II. Master Equation

The dynamics of protein level, as illustrated by Fig 1, left, is described by two components: the deterministic law that governs protein decay between bursts of gene expression; a probability transition law that governs the occurrence and size of individual gene-expression bursts. For the decay law we use a simple linear model d*x*/d*t* = −*γx*, where *γ* is the decay rate constant. The linear rate law implies that the protein level trajectories are piecewise exponential. The probability transition law is described by the burst kernel *B*(*x*|*y*), which gives the probability per unit time that a sufficiently large burst occurs which increases the protein level beyond the value *x*, provided that it currently stands at value *y*. In particular, *B*(*y*|*y*) gives the stochastic rate with which bursts (of any size) occur; the rate may or may not depend on the protein level *y*. The ratio *F* (*x*) = *B*(*x*|*y*)/*B*(*y*|*y*), where *x* > *y*, gives the upper-tail probability distribution function of post-burst level *x* conditioned on pre-burst level *y*.

In the simplest possible case of no feedback, the burst kernel is given by [1], [19]

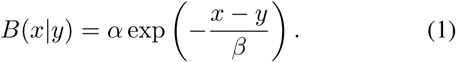

Equation (1) implies that (i) burst sizes are exponentially distributed with mean *β* and that (ii) they occur randomly in time with intensity (or burst frequency) *α* per unit time. We refer to kernel (1) as the constitutive kernel.

The probability distribution of protein level is obtained by formulating and solving an appropriate master equation. In discrete models of gene expression, master equations are systems of (typically infinitely many) ordinary differential equations [32]. In hybrid and piecewise deterministic models, such as that in Fig 1, left, master equations are described by (a limited number of) partial differential equations [33], which can also include a non-local integral term if discontinuous bursts are present in the dynamics [19]. In our case, the probability *p*(*x, t*) that protein is present at level *x* at time *t* satisfies a partial integro-differential equation [19], [28]

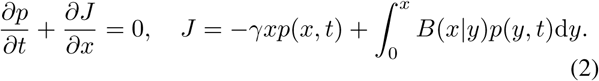

The first equation in (2) states the principle of probability conservation in differential form. The probability flux *J* includes a local flux −*γxp*(*x, t*) due to deterministic protein decay with rate constant *γ*. It also comprises a non-local integral flux due to bursts of protein synthesis featuring a burst kernel *B*(*x*|*y*).

The stationary probability density function (pdf) satisfies [28], [30]

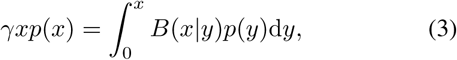

which is a homogeneous Volterra integral equation of the second kind. For the constitutive kernel (1) the stationary pdf is given by the gamma distribution [19]

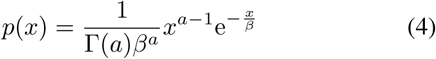

with shape *a* = *α*/*γ* (normalised burst frequency) and scale *β* (mean burst size). Elementary analysis of (4) shows that the gamma pdf is a unimodal distribution. Its mode is singular if its shape is less than one i.e. if the burst frequency *α* is less than the decay rate constant *γ*. Otherwise, the pdf possesses a regular peak at *x* = (*α*/*γ*−1)*β*. The mean of gamma distribution is equal to *αβ*/*γ* and its variance is equal *αβ*^2^/*γ*. The squared coefficient of variation, a widely used dimensionless measure of noise which is defined as the ratio of variance to mean squared, is therefore equal to the reciprocal burst frequency *γ*/*α*. Thus, in the absence of regulation, the steady-state distribution of a protein produced in bursts is unimodal, its mean is proportional to the burst frequency *α* and its coefficient of variation is inversely proportional to *α*. In the next sections, we will consider how the introduction of positive feedback affects these properties.

We shall consider two additional burst kernels, which correspond to two different regulation pathways, namely:

1. Feedback in burst frequency:

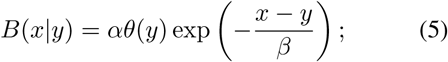
2. Feedback in burst size:

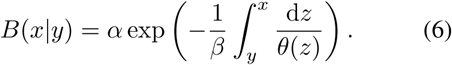

Feedback manifests itself in the burst kernels (5) and (6) through a response function *θ*(*x*). In the burst frequency feedback kernel (5), the response function alters the stochastic rate *B*(*y*|*y*) = *αθ*(*y*) with which bursts occur but not the probability distribution function *F* (*x*) = *B*(*x*|*y*)/*B*(*y*|*y*) = exp(−(*x* − *y*)/*β*) of the post-burst level. In the burst size feedback kernel (5), the stochastic burst rate *B*(*y*|*y*) = *α* remains unaltered but the response function modifies the post-burst level probability distribution function 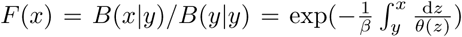. The function 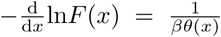 is referred to as the failure rate in reliability theory [34]. Hence, the response function directly modifies the rate of burst abortion.

If the response function is trivially given by *θ*(*x*) ≡ 1, then both (5) and (6) reduce to the constitutive kernel (1). In this paper, we shall consider a non-trivial example of an increasing response function *θ*(*x*), which corresponds to positive autoregulation. Specifically, we consider the non-cooperative Hill function

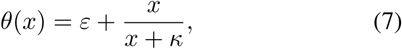

where *ε* is a (typically small) leakage parameter and *κ* gives a critical protein level, at which half-activation is achieved.

Without loss of generality we may assume that *β*= 1 and *γ* = 1. These choices mean that protein level is measured in units of the referential burst size *β* and time is measured in units of protein lifetime 1/*γ*. Having made these choices necessitates a minor reinterpretation of the other parameters *α* and *κ*. Specifically, *α* says how many bursts occur on average per protein lifetime and *κ* how many average-sized bursts amount to a critical level in terms of the regulatory feedback (7). The remaining parameter *ε*, having been non-dimensional to begin with, retains its original interpretation of leakage.

## III. Feedback In Burst Frequency

### A. Probability density and its moments

Assuming *β* = *γ* = 1, and inserting the burst-frequency kernel (5) into (3), we obtain

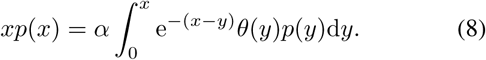

Multiplying (8) with e*^x^* yields

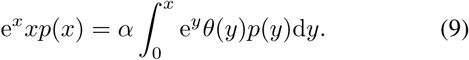

Differentiating (9) with respect to *x* gives

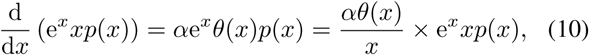

from which

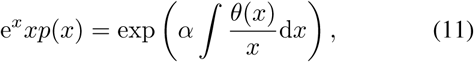

i.e.

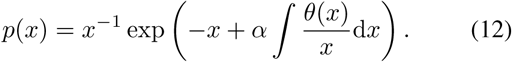

Note that the density function *p*(*x*) is not necessarily normalised. In general, any constant multiple of (12) is also a valid solution to (8).

The derivative of the density function (12) satisfies

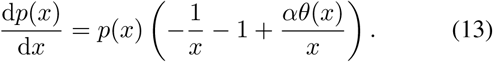

Critical points of the density satisfy d*p*(*x*)/d*x* = 0, which leads to

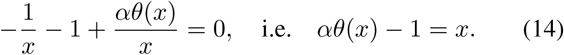

Elementary analysis of equation (14) with the noncooperative Hill function (7) implies that up to two critical points may exist.

In the specific case of non-cooperative Hill-type response (7), the density function *p*(*x*) is given explicitly by

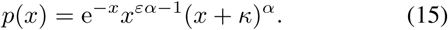

The *n*-the moment of the density function (15) is given by

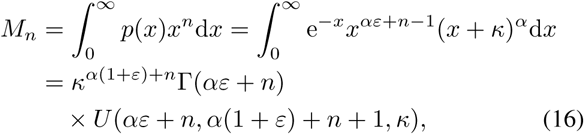

where Γ(*z*) is the gamma function and *U*(*a, b, z*) is the confluent hypergeometric function of the second kind [35].

### B. Parameter space

The pdf (15) can change from unimodal to bimodal (or back) in two ways: first, by developing (or losing) a singularity at the origin, which occurs if

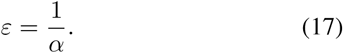

Additionally, modality may change if the critical-point equation (14) possesses a degenerate root.

The conditions for a degenerate root are

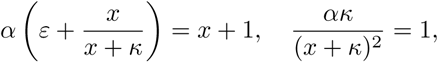

which, upon eliminating *x*, yield

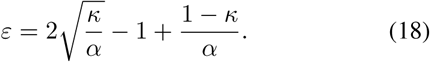

For a fixed value of *κ* > 0, conditions (17) and (18) represent two curves in the (*ε,α*)-parameter space delineating the region of bimodal behaviour (Fig 2).

**Fig. 2.**
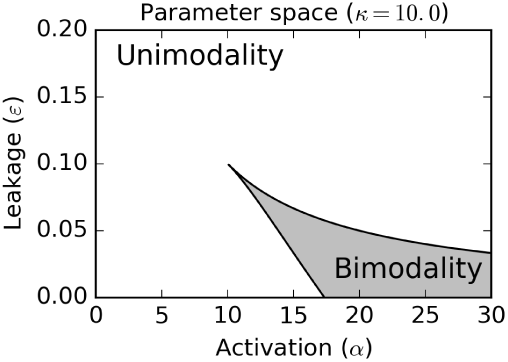
Parameter space of the model for feedback in burst frequency with dimensionless critical concentration threshold fixed to *κ* = 10.

## IV. Feedback In Burst Size

Again assuming *β* = *γ* = 1, and inserting the burst-size regulation kernel (6) into (3), we obtain

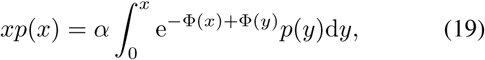

where

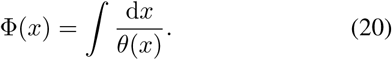

Multiplying (19) with e^Φ(*x*)^, we obtain

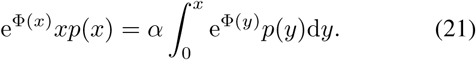

Differentiation of (21) yields

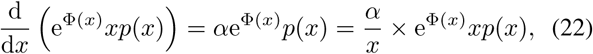

so that

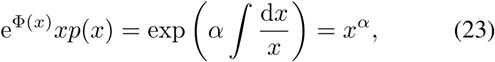

i.e.

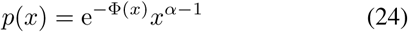

is the desired solution.

The derivative of the density function (24) satisfies

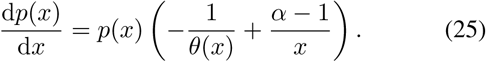

Critical points satisfy

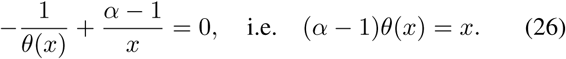

Elementary analysis of equation (26) with the noncooperative Hill function (7) implies that more than one critical point cannot exist. Therefore, the probability density function remains unimodal throughout the parameter space.

For non-cooperative Hill-type response (7), the density function *p*(*x*) in (24) takes the explicit form of

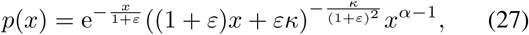

and

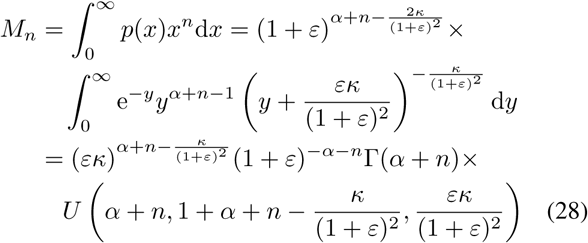

gives the *n*-the moment of the density function (15).

## V. Discrete Models

In the previous sections, we derived protein probability density functions for hybrid models of feedback in burst frequency and size. Here we compare the hybrid analysis with fine-grained discrete chemical-kinetics models. The discrete models of burst size and frequency feedback include three species: the inactive state (I), the active state (A), and the protein (P). Burst-like limit is achieved if the active state has short holding times but drives rapid protein production.

In case of frequency regulation, the rate of activation depends on protein amount. The reaction system is given by

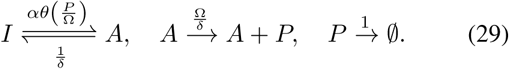

The response function *θ*(*x*) is given by (7). The parameter *δ* compares the bursting and protein turnover timescales. Bursting limit is achieved by taking *δ* ≪ 1. The mean number of protein produced per burst plays the role of a system-size parameter and is denoted by Ω. It is expected, and numerically verified in the next section, that in the limits of Ω → ∞ and *δ* → 0 the protein concentration defined by *x* = *P/*Ω satisfies the hybrid model (2) and (5) for feedback in burst frequency (cf. [21], [36]).

The reaction system for feedback in burst size reads

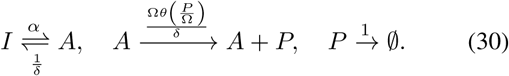

Again, it is expected that the hybrid model for feedback in burst size given by (2) and (6) is achieved from (30) in the limit of Ω → ∞ and *δ* → 0.

## VI. Results

In this section we apply the theoretical analysis of Sections III–V to gain an overview of the model behaviour. In Figs 3 and 4 we show (i) the steady-state protein mean and squared coefficient of variation (CV^2^) as function of referential burst frequency *α* and (ii) the steady-state protein distribution for selected values of *α*. The CV is defined the ratio of the standard deviation to the mean value. Biologically, an increase in the referential burst frequency can be interpreted as resulting from the turning on of an upstream activatory pathway. Therefore we shall refer to *α* as to the activation parameter and relate the magnitude of *α* to the level of upstream transcriptional activation of the gene. We use a selection of values for the leakage parameter *ε* and fix the value of the critical concentration to *κ* = 10.

**Fig. 3.**
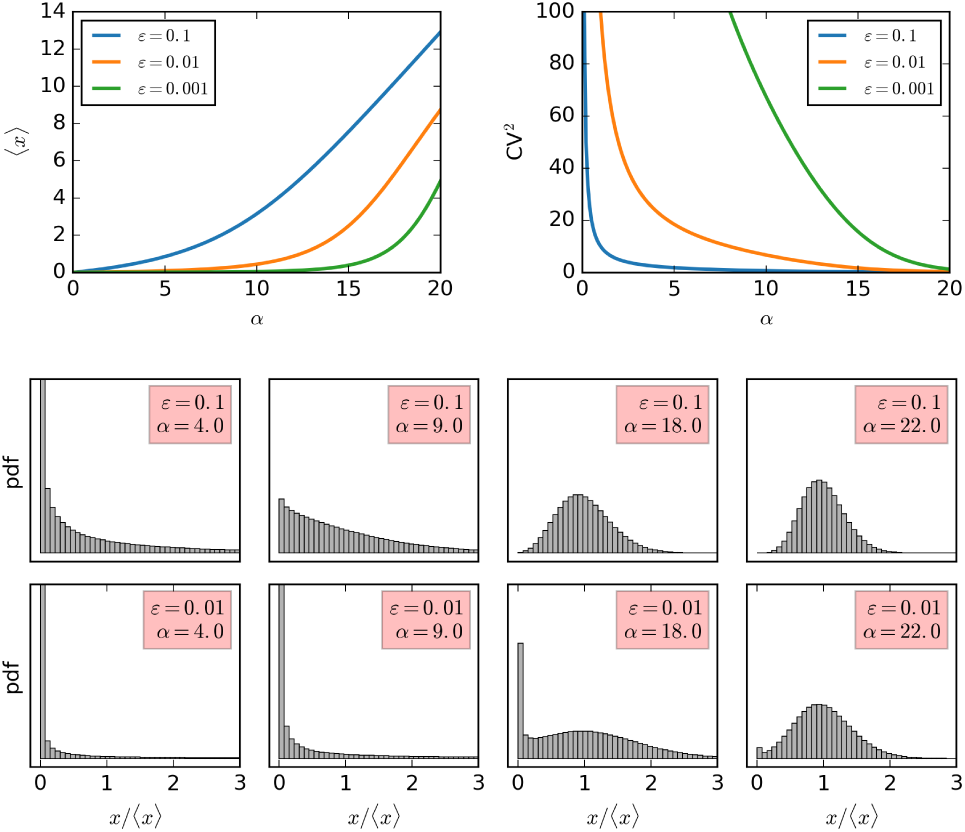
Steady-state protein moments and probability density functions in response to gene activation (*α*) subject to noncooperative positive feedback in burst frequency. Parameters: the critical protein level is set to *κ*= 10 throughout; the activation parameter *α* is varied from low to high values; a selection of leakage values *ε* is considered.

In the case of noncooperative positive feedback in burst frequency (Fig 3), the protein mean is an increasing function, and the protein noise, i.e. the CV^2^, is an decreasing function of the activation parameter *α*. For very low values of leakage, protein exhibits low means even at high activation levels, and is extremely noisy at low activation levels. The protein probability density function (pdf) is bimodal for small values of leakage parameter and sufficiently large activation levels. The trivial lower mode *x* = 0 corresponds to the absence of protein. The pdf is unbounded at *x* = 0; in order to correctly appreciate how much probability mass is concentrated near the singularity at *x* = 0, the probability density function is binned in Fig 3.

In the case of feedback in burst size (Fig 4), the protein mean also increases with the activation parameter *α*. In contrast to burst-frequency regulation, there is a lower bound 〈*x*〉 > *α* − *κ* for the protein mean which is valid even as the leakage parameter *ε* tends to zero. We can say that as *α* exceeds *κ*, the protein becomes activated regardless of the value of leakage. The protein noise decreases monotonically with in response to activation only if the leakage *ε* is relatively large. If the leakage is sufficiently small, the noise exhibits a transient spike shortly before the protein becomes activated at *α* = *κ*. The protein distribution is unimodal across the parameter space.

**Fig. 4.**
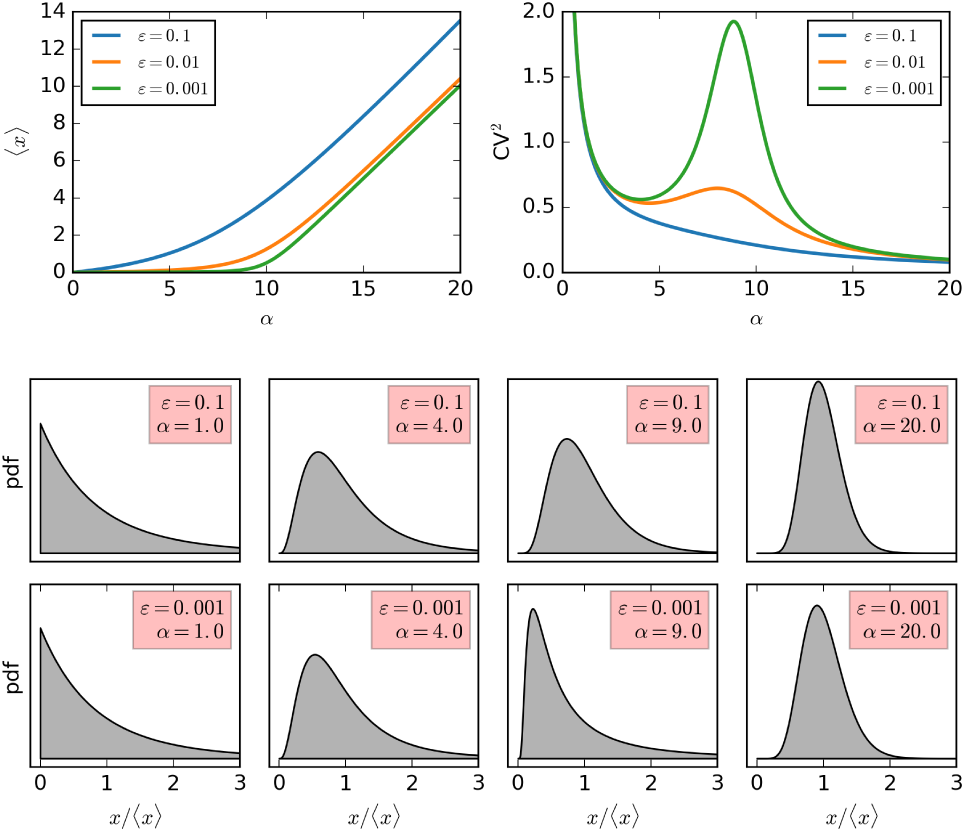
Steady-state protein moments and probability density functions in response to gene activation (*α*) subject to noncooperative positive feedback in burst size. Parameters: the critical protein level is set to *κ* = 10 throughout; the activation parameter is varied from low to high values; a selection of leakage values *ε* is considered.

In Figs 3 and 4, the probability density function (pdf) was obtained by normalising the density funcions (15) and (27) with the corresponding zero-th moments *M*_0_ (16) and (28). The formulae

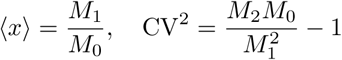

were used to calculate the mean and the CV^2^ from the moments (16) and (28).

We compare the continuous modelling framework to the results of kinetic Monte Carlo simulations of full discrete models in Fig 5. In addition to the parameters of the continuous framework, the critical concentration *κ*, leakiness *ε*, and gene activation *α*, the discrete formulations include two additional parameters. The system size parameter Ω gives the number *P* of protein molecules that correspond to the unit of protein concentration *x*. Continuity in protein level is achieved in the limit of Ω → ∞. The parameter *δ* compares the length of bursting and protein turnover timescale; bursts become instantaneous in the limit of *δ* → 0. The protein copy-number histograms obtained by stochastic simulation are compared to the transformed probability density functions

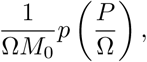

where *p*(*x*) is the protein pdf in the presence of feedback in burst freqency (15) or burst size (27); the normalisation constant *M*_0_ is equal to the zero-th moment (see (16) and (28)).

**Fig. 5.**
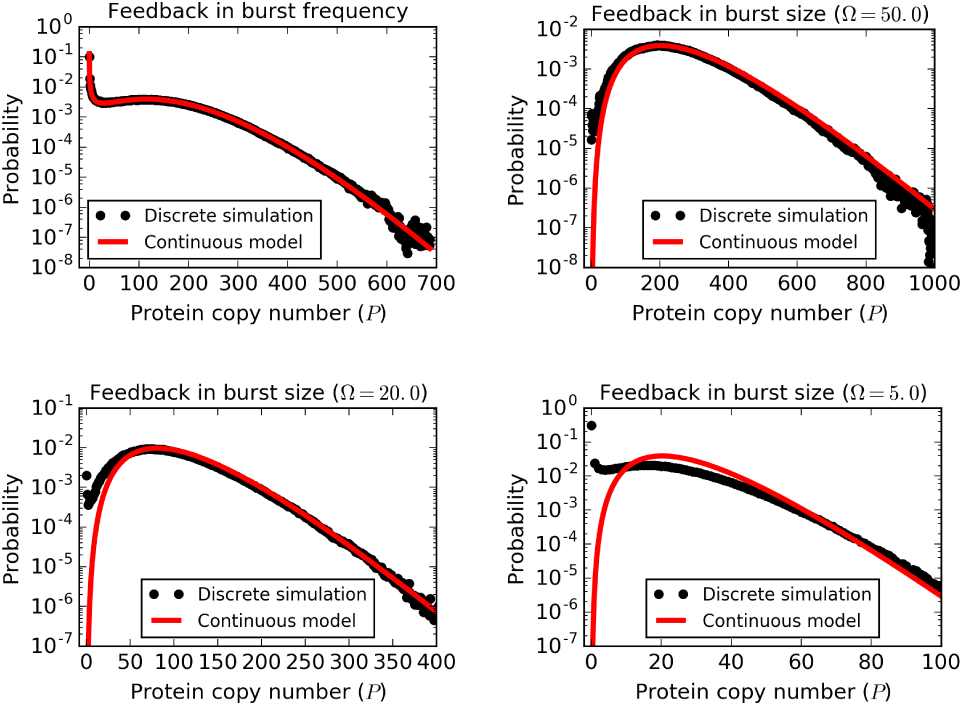
Steady-state protein copy number distributions for feedback in burst frequency (top left panel) and feedback in burst size (other panels). Simulations of full discrete models (29) and (30), as discrete black markers, are compared to explicit continuous pdfs (15) and (27), as solid red lines. The parameters are *κ* = 10, *α* = 18, *ε* = 10^−2^, Ω = 20, δ = 10^−3^ for burst frequency regulation and *κ* = 10, *α* = 15, *ε* = 10^−3^, *δ* = 10^−3^, Ω ∈ {5, 20, 50} for burst size regulation. The simulation result of each panel is based on 10^7^ direct method iterations of the appropriate discrete stochastic model performed in StochPy [37].

The discrete model for feedback in burst frequency is in an excellent agreement with the hybrid framework (Fig 5, top left panel). For feedback in burst size, the two modelling approaches agree well if the system size is sufficiently large (Fig 5, top right panel). However, at lower system sizes, the discrete model for burst size feedback leads to higher probabilities of having zero or few protein molecule copy numbers (Fig 5, bottom panels). This can ultimately lead to bimodal distributions, with one mode corresponding to the absence of protein and the other mode corresponding to the up-regulated regime (Fig 5). We conclude that bimodality can arise in case of feedback in burst size due to low protein copy number effects.

Let us try to provide an intuitive explanation for the presence and absence of bimodality in protein subject to positive nooncooperative feedback in burst frequency or size. Bimodality occurs as a result of pseudo-extinction. Protein level is normally supported by the feedback loop, but can drop to near zero values due to random fluctuations, at which the feedback is interrupted and any production is due to leakiness. In case of feedback in burst frequency, this means that there is a large waiting time for the next burst which rescues protein from pseudo-extinction state. The long holding time in the pseudo-extinction state drives the emergence of bimodal protein distributions. In case of feedback in burst size, bursts retain their frequency but are smaller if protein level detours into low values. Since every small burst makes the next burst likely to be a little larger, the possibility of pseudo-extinction is averted in burst-size regulation. Nevertheless, if low copy numbers of protein are considered, very small bursts will simply have the size of zero. This means that the waiting time until the next *nonzero* burst from the pseudo-extinction state can actually be very large even in case of feedback in burst size if low protein copy numbers are taken into account. Therefore, bimodality can emerge for low copy number protein subject to feedback in burst size. Nevertheless, at reasonably large copy numbers, feedback in burst size is less conducive to bimodality than feedback in burst frequency, as suggested by the detailed analysis of the hybrid model.

## VII. Conclusions

We considered the effects of noncooperative positive feedback in burst size or burst frequency on steady-state protein distributions using a hybrid gene-expression model. Feedback in burst frequency was shown to support bimodal protein distribution without cooperativity, whereas feedback in burst size typically does not, except for low protein copy number driven bimodality. The two types of feedback led to strikingly different values of protein mean and protein noise (coefficient of variation), in particular in small-leakage scenarios. Our analysis was based on exact solutions to the master equation and stochastic simulation of a fine-grained discrete chemical kinetics model. The current results contribute to the understanding of the distinction between different types of gene-expression regulation. The methodologies used to obtain these results are likely to be useful in other contexts also.

